# Limitations of Current Machine-Learning Models in Predicting Enzymatic Functions for Uncharacterized Proteins

**DOI:** 10.1101/2024.07.01.601547

**Authors:** Valérie de Crécy-Lagard, Raquel Dias, Nick Sexson, Iddo Friedberg, Yifeng Yuan, Manal A. Swairjo

## Abstract

Thirty to seventy percent of proteins in any given genome have no assigned function and have been labeled as the protein “unknome”. This large knowledge shortfall is one of the final frontiers of biology. Machine-Learning (ML) approaches are enticing, with early successes demonstrating the ability to propagate functional knowledge from experimentally characterized proteins. An open question is the ability of machine-learning approaches to predict enzymatic functions unseen in the training sets. Using a set of *Escherichia coli* unknowns, we evaluated the current state-of-the-art machine-learning approaches and found that these methods currently lack the ability to integrate scientific reasoning into their prediction algorithms. While human annotators can leverage the plethora of genomic data in making plausible predictions into the unknown, current ML methods not only fail to make novel predictions but also make basic logic errors in their predictions. This underscores the need to include assessments of prediction uncertainty in model output and to test for ‘hallucinations’ (logic failures) as a part of model evaluation. Explainable AI (XAI) analysis can be used to identify indicators of prediction errors, potentially identifying the most relevant data to include in the next generation of computational models.

**Article Summary:** Many proteins in any genome, ranging from 30% to 70% of the genome, lack an assigned function. This knowledge gap limits the full use of the vast available genomic data. Machine learning has shown promise in transferring functional knowledge within isofunctional families, but it largely fails to predict novel functions not seen in its training data. Understanding these failures can guide the development of better machine-learning methods to help experts make accurate functional predictions for uncharacterized proteins.

## Introduction

Determining protein function is not an easy task, and thirty years after the first bacterial genome was sequenced, the functional annotation status of the proteome of most species is far from being accurate or complete, even for model organisms (Ghatak et al., 2019; Wood et al., 2019; de Crécy-lagard et al., 2022; Rocha et al., 2023). Experimental validation of protein function is a painstaking process, and with the explosion of whole genome sequences (Kyrpides, 1999; Peterson et al., 2025; Torres et al., 2025), the gap between experimentally validated functions and those predicted through computational methods continues to widen. In UniprotKB (Bateman et al., 2023), the most widely used protein function database (Ramola et al., 2022), the estimates are that less than 0.5% to 15% of proteins have been linked to experimental data(Škunca et al., 2017).

The process of functional annotation of protein entries in databases starts with capturing information in the literature by biocurators(International Society for Biocuration, 2018). This process links experimental characterizations of specific proteins in specific organisms to controlled vocabularies that describe validated functions, such as the Gene Ontology (GO)(Gene Ontology Consortium et al., 2023), the IUPAC Enzyme Commission (EC) numbers (https://iubmb.qmul.ac.uk/enzyme/), or biochemical reaction descriptors [e.g. Rhea(Bansal et al., 2022)]. Text-mining tools have accelerated the flow of information captured(Soldatos et al., 2015; Poux et al., 2017; Wei et al., 2019). However, this step remains a major bottleneck in the annotation workflow, which can result in mislabeling proteins as “unknown” when a function has been reported in the literature (see type 1 error in Table 1 and Fig. 1).

**Figure 1.**
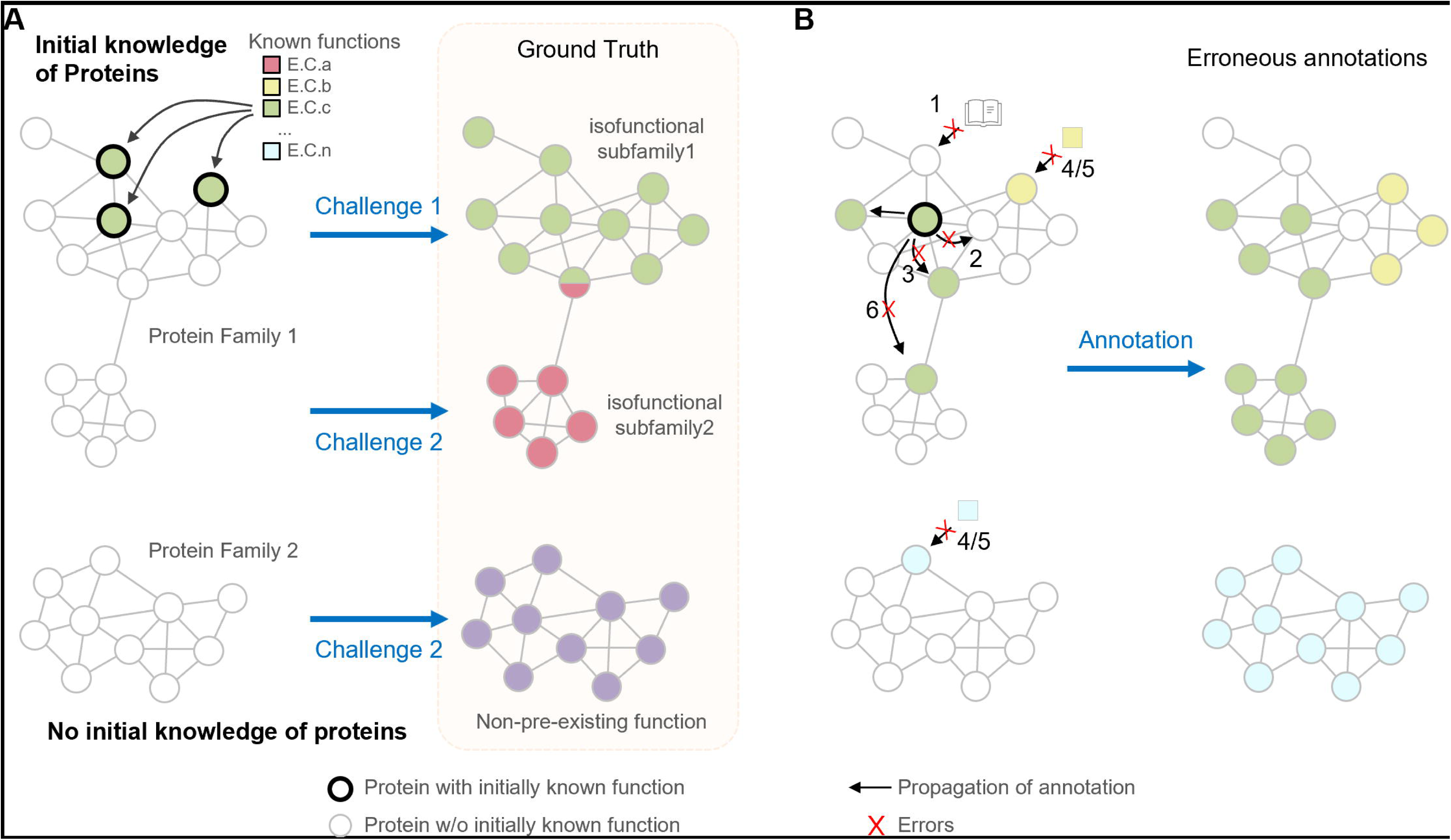
The major two types of challenges in accurate protein functional annotation. (A) Challenge 1: to propagate the existing knowledge to the correct set of unannotated proteins. Challenge 2: to annotate proteins in a family with none of the members linked to any initial known information. Each circle represents a protein in a family with (black) or without (gray) initially known function. The edges that connect two circles represent protein similarity above a preset threshold. The squares represent known functions with E.C.a to E.C.n as examples. (B) The errors during propagation of annotation that could lead to erroneous results. The numbering of errors (red crosses) corresponds to the error types described in Table 1.

**Table 1:** Main types of errors that lead to erroneous functional annotations of proteins. Error types 1-6 error types correspond to the errors numbered in Figure 1.

| **Error Types** |  |  |
| --- | --- | --- |
| ***False unknowns*** | **Annotated as unknown or general but function is known and published** |  |
|  | ***Details*** | ***Example*** |
| 1 Failure to capture the literature | Not captured in any database or captured in some databases but not others. Also annotated as vague when precise annotation is known. | CT_611 is captured as folylpolyglutamate synthase in KEGG (ctr:CT_611) but this annotation is still not in Uniprot as of April 2024 (O84617). See also examples in (Price and Arkin, 2024). |
| 2 Naming issues<br>Propagation failures | Inconsistent naming of the same entities. Captured for one member of a family but not propagated to others. | See GroEL in (Lockwood et al., 2019). The MptE protein that encodes 6-hydroxymethyl-7,8-dihydropterin pyrophosphokinase in most Archaea (De Crécy-Lagard et al., 2012) is captured in BioCyc for *Methanocaldococcus jannaschii* DSM 2661 (https://biocyc.org/gene?orgid=MJ&id=MJ_RS08700-MONOMER) but not propagated to any archaeal homolog. |
| ***False knowns*** | **Annotated as precise but wrong (should be general or another annotation) or incomplete** |  |
|  | ***Details*** | ***Example*** |
| 3 Multiple functions | Protein has multiple functions because of fusions moonlighting or promiscuity and only one of the functions is captured. | Very common few fusions have been linked to the multiple encoded functions. For example, A5I019 has two functions QueD and PTPS-III only one is called in Uniprot (de Crécy-Lagard, 2014). See also examples in (Price and Arkin, 2024). |
| 4 Curation mistake | 1) The data was incorrectly captured by biocurator; 2) Functional annotations may become outdated. | Ureidoglycolate lyase (Percudani et al., 2013). |
| 5 Experimental mistake | Finding has been refuted by other studies: 1) the published data is inconclusive; 2) Inconsistency occurs when different databases or resources provide conflicting functional annotations for the same protein. | DUF34 family was annotated as GTP cyclohydrolase 1B (Reed et al., 2021). |
| 6 Over-annotation of paralogs | Annotation wrongly propagated to non-isofunctional paralogous groups. | See examples in (Schnoes et al., 2009; Zallot et al., 2016; Rembeza and Engqvist, 2021). |

Using the principle of sequence similarity, also known as homology transfer, putative functions are assigned to proteins in newly sequenced genomes as a part of the genome annotation process (Seemann, 2014; Thibaud-Nissen et al., 2016; Olson et al., 2023). Indeed, most functions of proteins in UniProt have been inferred computationally based on sequence similarity (Bateman et al., 2023). The methods used over the past 30 years to automatically propagate functional annotations have been extensively reviewed (Storm and Sonnhammer, 2002; Friedberg, 2006; Lee et al., 2007; Satish Kumar et al., 2007; Devoid et al., 2013). This seemingly simple task can become quite difficult (Challenge 1 in Fig. 1), since relying on sequence similarity alone can result in significant annotation errors (Schnoes et al., 2009; Rembeza and Engqvist, 2021). One prevalent error type affecting protein families (Rembeza and Engqvist, 2021) is in the misannotation of paralogs (Schnoes et al., 2009; Zallot et al., 2016; Rembeza and Engqvist, 2021) caused by the inherent functional complexity of enzyme families (Glasner et al., 2006; Gerlt et al., 2012) (see type 6 error in Table 1 and Fig. 1). Functional diversification through duplication and divergence (Todd et al., 2001; Das et al., 2015; Bordin et al., 2021) results in proteins with high degrees of similarity having different functional roles, creating non-isofunctional paralogous groups. Indeed, even within a single protein family, a difference in a few amino acids can affect substrate binding and/or catalysis, effectively changing the function (Roy et al., 2009; Ribeiro et al., 2020; Precord et al., 2023). Most current genome annotation pipelines annotate paralogs without considering the potential for functional divergence, leading to incorrect annotations for up to 80% of family members, with these types of errors increasing over time (Schnoes et al., 2009; Rembeza and Engqvist, 2021).

Methods for inferring function that incorporate additional information such as phylogeny, active site analyses, metabolic reconstruction, and sequence similarity networks (SSNs) combined with gene neighborhood or co-expression information can better separate non-isofunctional subfamilies (Zallot et al., 2016, 2021; Ribeiro et al., 2023) (Fig. 2). When Hidden Markov Models (HMMs) or signature motifs are generated to annotate the subfamilies, they can be integrated into annotation pipelines such as RefSeq(Li et al., 2021) or incorporated into rules such as the as the Unified Rule (UniRule) used by UniprotKB (MacDougall et al., 2020) additional curation, which has not kept pace with the exponential increase in sequenced genomes. In addition to mis-assignments among paralogs increasing with evolutionary distance, distinguishing between sub- and neo-functionalization can be problematic (Birchler, 2025).

**Figure 2.**
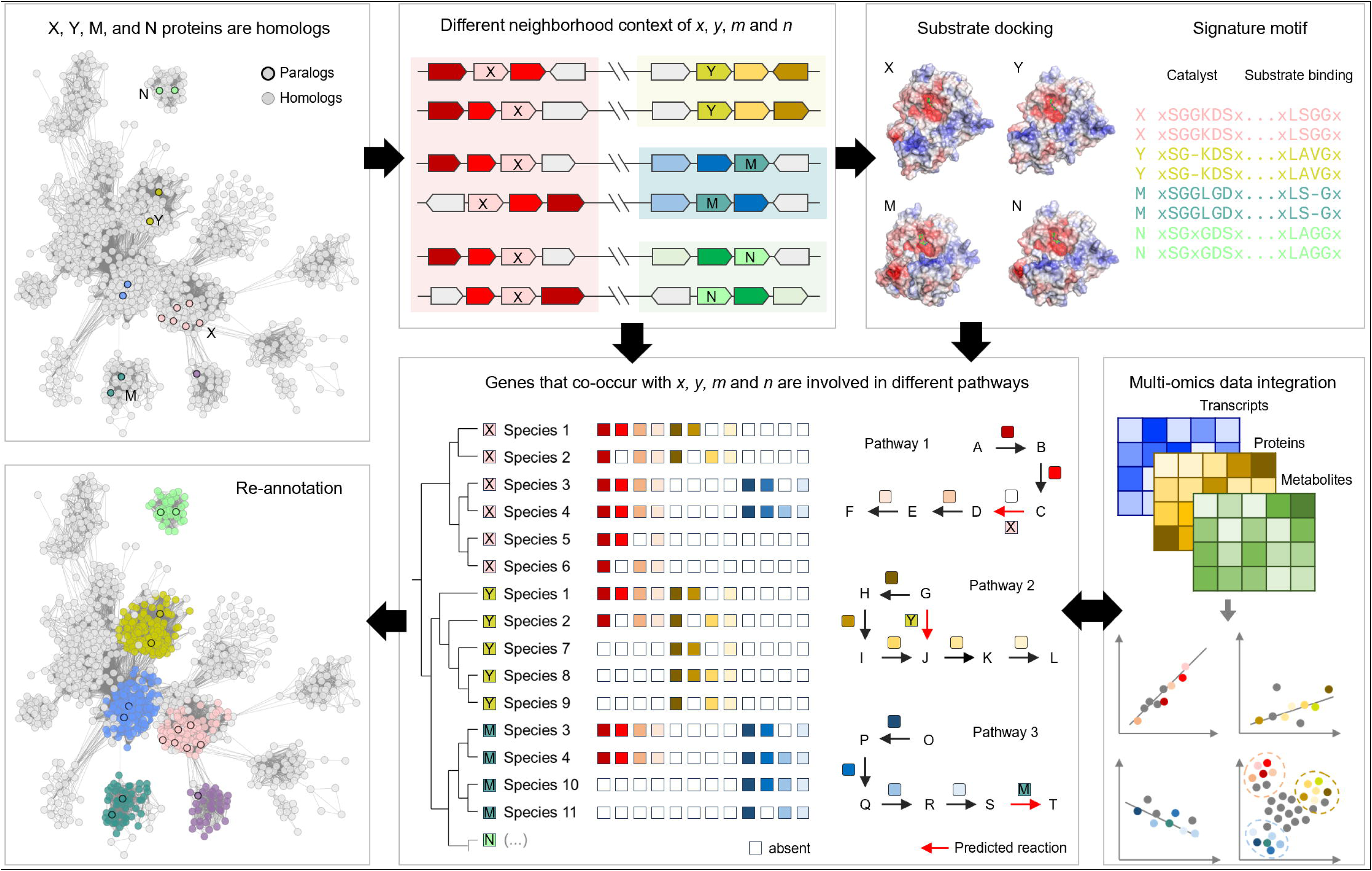
Computational workflow used to separate non-isofunctional paralogs in a protein superfamily. Sequence similarity networks of protein families can separate paralogs by integrating different types of information, Arrows indicate the flow of information. The final-colored network shows the paralog families that could be separated.

A significant challenge is to predict the functions of the proteins that have not previously been functionally characterized (Fig. 1A). Estimates suggest that 30% to 70% of proteins in any given genome are in this “unknown” category (Ghatak et al., 2019; Wood et al., 2019; Lobb et al., 2020; Sajid et al., 2023). Functionally characterizing these unknown families is challenging because of the openness of the prediction space (Hanson et al., 2009; Niehaus et al., 2015). Some of these “unknowns” are likely to fill functional roles not yet linked to a specific gene (Karp, 2004; Lespinet and Labedan, 2006; Chen and Vitkup, 2007; Niehaus et al., 2015). Others are likely to be non-orthologous replacements (or convergent evolution), where different (structurally diverse) protein families overlap in functional roles (Ribeiro et al., 2023). Finally, we expect some novel biological functions and predicting these require extrapolation beyond the current knowledge base. As we sequence more organisms from more previously unexplored ecological niches, we expect to encounter proteins with novel functions.

In summary, the annotation of a new genome faces two challenges: 1) to identify genes whose likely function can be inferred by propagating known annotations; 2) to identify genes with potentially novel functions and make predictions. This is a thorny path with many different types of errors that can be made (Fig. 1 B and Table 1). Computational approaches, however innovative, cannot prove a particular protein function; experimental validation is still necessary. Proof of function is not necessarily straightforward, as demonstrating a protein capable of participating in a chemical reaction is not proof that the protein has the responsibility of that function in the cellular context where it is identified (García-Contreras et al., 2012; Copley, 2015). For proteins with available experimental evidence of function in one context, the computational propagation of the assigned functional role is the logical next step (Lee et al., 2007), although as described above, similarity does not guarantee functional conservation. For unknown proteins, the increasing amounts of genome-wide/transcriptome-wide/metabolome-wide experimental data can be used in combination with structural knowledge to develop predictions that can be tested experimentally (Hanson et al., 2009). Scientists are increasingly successful in these types of endeavors in a range of systems (Dembech et al., 2023; Rodríguez del Río et al., 2023), and there is tremendous hope that AI methods will accelerate the pace at which predictions can be made.

For challenge 1, there has been a recent explosion of publications reporting the use of pre-trained Protein Language Models (PLM) to predict protein functions (Ardern et al., 2023; Ayres et al., 2023; Derry and Altman, 2023; Durairaj et al., 2023; Kim et al., 2023; Prabakaran and Bromberg, 2023; Rodríguez del Río et al., 2023; Hwang et al., 2024) (Fig. 3). Models have been developed to link protein sequences directly to vocabularies such as GO terms (Radivojac et al., 2013; Jiang et al., 2016; Zhou et al., 2019) or EC (Enzyme Commission Classification) numbers (Hamamsy et al., 2023). EC numbers are a set of 4 numbers that are hierarchical with the first number the most general classification and the last number the most specific, and whose purpose is to standardize the description of enzymatic activities. PLMs have been reported to accurately predict the first two digits of the EC number but not the last two (Sanderson et al., 2023). GloEC (Huang et al., 2024) and MAPred (Rong et al., 2024) are two recent PLM-based EC prediction tools (Fig. 3) that achieve precision and recall metrics combined into macro-F1 or regular F1 scores ranging from 10% to 80%, depending on the test set [see Tables 2-4 of (Huang et al., 2024) and Table 1 of (Rong et al., 2024) respectively, summarized in Fig 4]. The F1 score is the harmonic mean of precision and recall. Recall is the proportion of true positives correctly identified out of all actual positives, while precision is the proportion of true positives among all predicted positives. A “positive” refers to an instance that actually belongs to the class of interest (here, the correct EC number). The macro-F1 score is calculated by computing the F1 score for each EC number category independently and then averaging these scores across EC numbers. When training data contains well-characterized proteins with well-described structures (e.g., cofactor-237 set) or well-separated functional groups (e.g., phosphorylase set), F1 scores range from 60% to 75% for the best methods (Fig. 4A). Once the structure of the training data is more complex, and the algorithm must distinguish among similar enzymes with different functions, such as the carbohydrate dataset comprised of ∼350 glycosyl hydrolases, F1 scores fall sharply, ranging from 6% to 30% (Fig. 4A and B). When using the Price test dataset, which comprises 100 enzymes that were annotated using high-throughput phenotypic data and are not included in any training sets(Price and Arkin, 2024), the best performance was a 50% F1 score with the MAPred pipeline (Fig. 4B). As recently discussed by the Bromberg group (Prabakaran and Bromberg, 2023)PLM-based methods are not designed to discover novelty (Challenge 2), as they rely on training data derived from existing knowledge.

**Figure 3.**
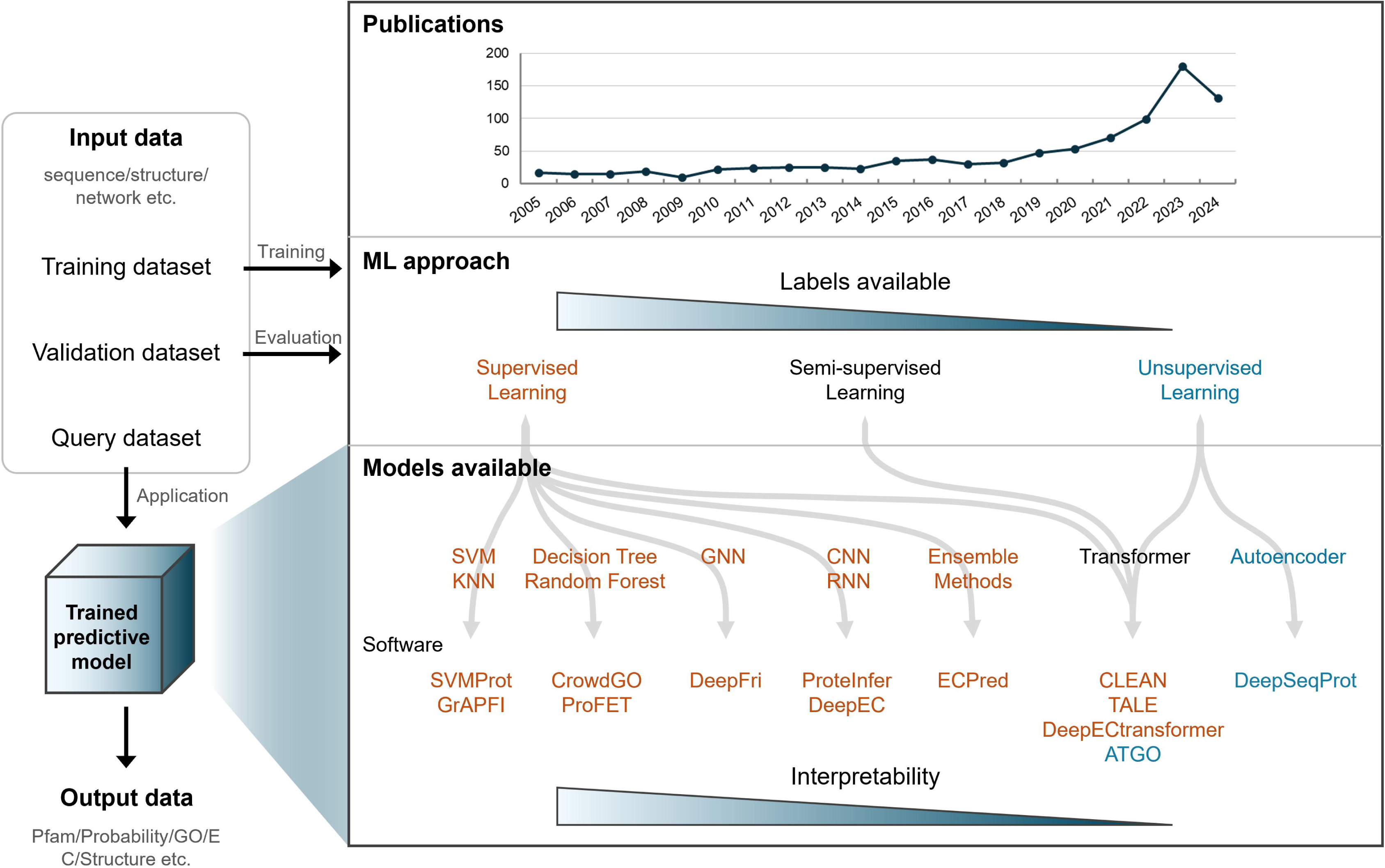
Workflow for functional annotation of proteins using machine learning. General steps are depicted: input data collection (training, validation, and query datasets), options for machine learning approach, model selection, training, and evaluation, and output data. Supervised, semi-supervised, and unsupervised learning approaches with examples of their respective modeling architectures and software, are colored differently. Blue scale bars represent the availability of labeled data (top) and the level of interpretability (bottom) of machine learning approaches and models available, respectively. The figure illustrates common associations between model architectures and machine learning approaches. However, it should be noted that models such as GNNs, CNNs, RNNs, and Transformers can be applied across supervised, semi-supervised, and unsupervised learning contexts. The total number of publications in each year was collected on June 10, 2024 from PubMed (pubmed.ncbi.nlm.nih.gov) using the search criteria: ‘function prediction protein language model’ in full data.

**Figure 4.**
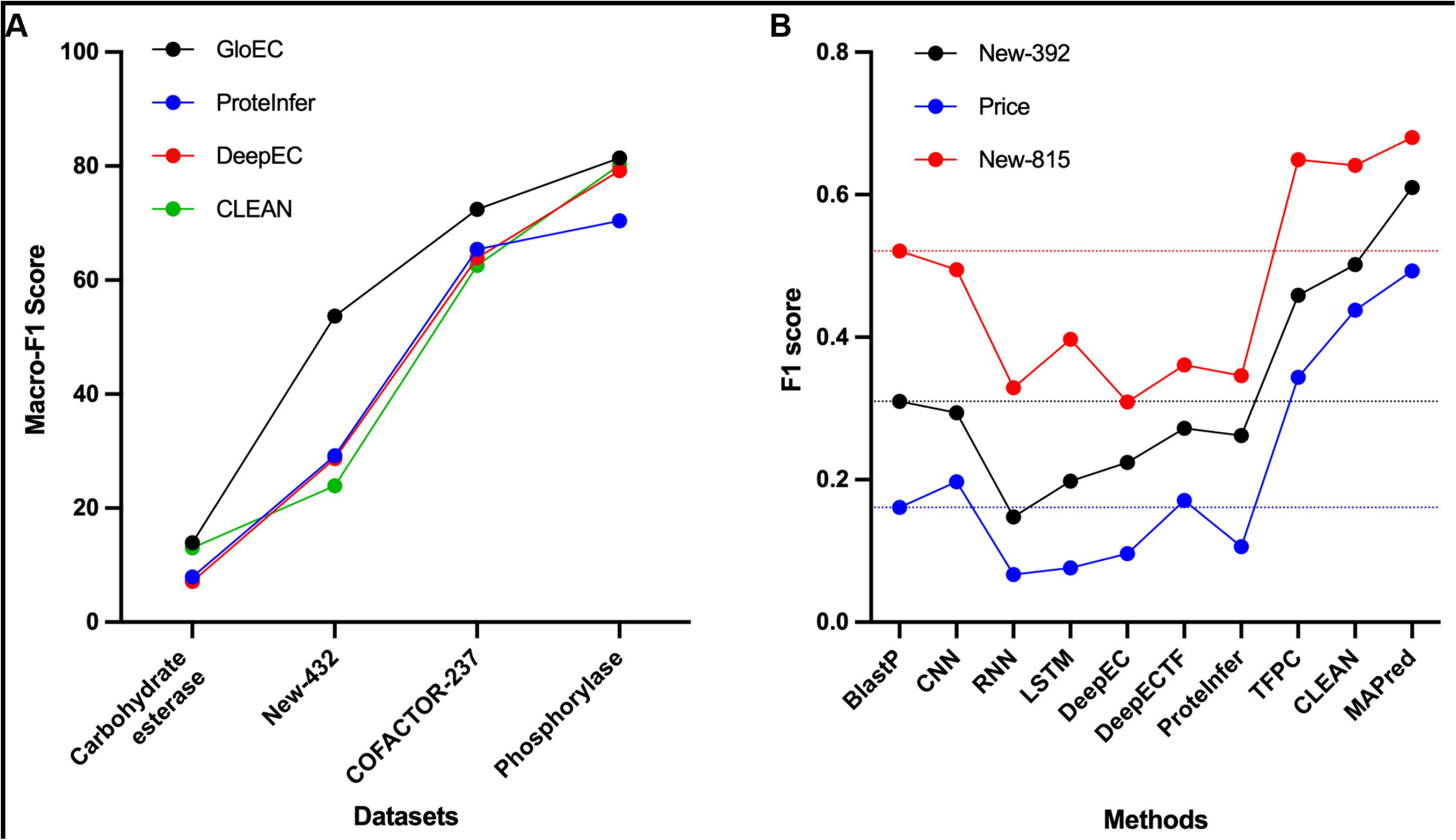
Comparison of different machine-learning methods to predict EC numbers. (A) Comparison of macro-F1 scores generated using four methods (GloEC, Protein Infer, DeepEC, and CLEAN) on four datasets (Carbohydrate esterase, New-432, Cofactor-237, and phosphorylase)(data extracted from (Huang et al., 2024)); (B) Comparison of F1-scores generated using eight methods including BlastP, DeepECTF, DeepEC, and CLEAN on three datasets (New-815, New-392 and Price)(data extracted from (Rong et al., 2024)). Dotted lines mark the BlastP scores. See initial publications for details and references for methods and datasets.

**Table 2.**
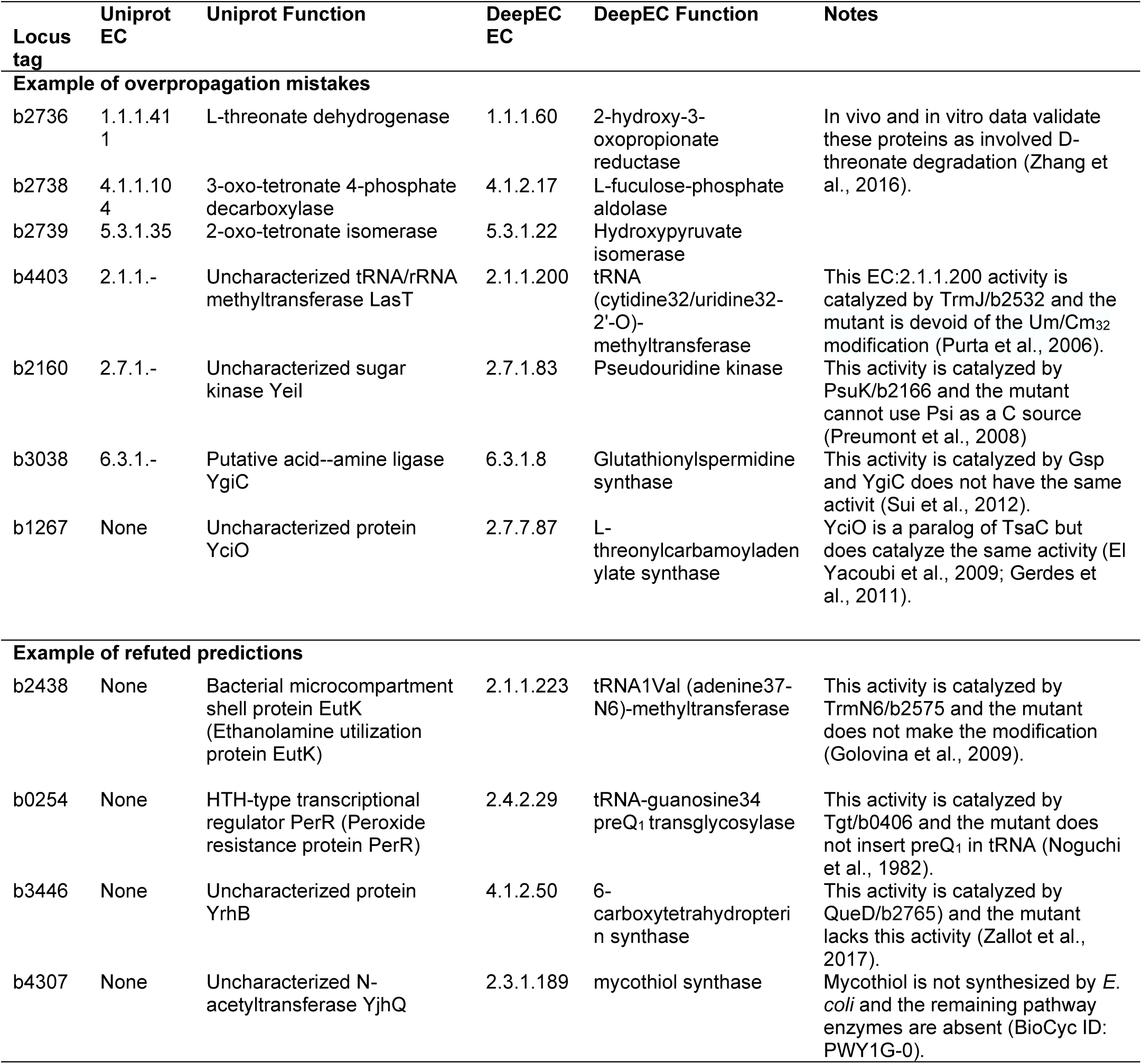
Example where experimental evidence contradicts the DeepECTF predictions.

Several independent studies agree that the number of proteins in the model gram-negative *Escherichia coli* K12 not linked to precise molecular and/or biological functions is between 1,200 and 1,400, or a quarter of the encoded proteins (Ghatak et al., 2019). Some of these proteins are likely responsible for functions that have been described in *E. coli* but never linked to a gene [called orphan enzymes (Karp, 2004; Danchin et al., 2018; Kobras et al., 2021)] or functions that might have been described in another organism but could also be present in *E. coli.* What percentage of these currently unannotated proteins represent novel functions is difficult to determine. A recent study labeled over 450 of the unknowns in the model bacteria *Escherichia coli* with substrate-level specificity ECs (all 4 numbers) using a supervised ML-based DeepECTransformer (DeepECTF) platform and validated three of these predictions *in vitro* (Kim et al., 2023). If corroborated, this report could be a real breakthrough in predictive modeling by linking true unknowns with their function, a process that usually takes years (Hanson et al., 2009; Niehaus et al., 2015). We designed a study to compare human curation to the reported DeepECTF predictions and examined in detail the potential of this new approach.

## Methods

### Comparing human curation with DeepECTF predictions

453 protein predictions from DeepECTF were grouped into different categories with confidence scores based on evidence assembled from human curation. Concordant predictions for all 4 EC numbers were given a confidence score of 2. These were of two types. Proteins with the identical functional annotation already present in UniProt (with or without an EC number), and proteins with functions not captured by UniProt. Discordant predictions were cases where: 1) another protein was known to perform the same function and experimental data showed that there were no redundancies; 2) the predicted function was known to be absent in *E. coli;* 3) the published literature or comments in UniProt provided evidence for a different function; were given a confidence score of 0. When proteins were members of families with many paralogous subgroups but no literature disproving the prediction was found and in other cases of uncertain calls, a confidence score of 1 was given.

### Bioinformatic analyses and data mining

The UniProt (www.uniprot.org)(Bateman et al., 2023), InterPro (www.ebi.ac.uk/interpro/)(Paysan-Lafosse et al., 2023), NCBI (www.ncbi.nlm.nih.gov)(NCBI Resource Coordinators, 2016), BioCyc (www.biocyc.org)(Karp et al., 2019) and KEGG (Kanehisa et al., 2021) knowledgebases were used to gather information on the 453 proteins. PaperBlast was used to query literature (https://papers.genomics.lbl.gov/cgi-bin/litSearch.cgi)(Price and Arkin, 2017). Open AI ChatGPT 4.o was used to extract data from Tables in publications and generate figures (https://chatgpt.com/) on August 20 using the prompt: “Extract data from Table X in attached pdf file”.

### Structural and phylogenetic analyses

Crystal structures were retrieved from the Protein Data Bank (www.rcsb.org)(Burley et al., 2022) and visualized using Pymol (www.pymol.org)(The PyMOL Molecular Graphics System, Version 1.8, Schrödinger, LLC). Structure-based multi-sequence alignment derived from available crystal structures and AlphaFold structural models (https://alphafold.ebi.ac.uk/) were generated using PROMALS3D (prodata.swmed.edu/promals3d/promals3d.php)(Pei et al., 2008) and Espript (<u>espript.ibcp.fr/ESPript/ESPript)</u>. The percent conservation scores were calculated using the program AL2CO (http://prodata.swmed.edu/al2co/al2co.php)(Pei and Grishin, 2001) using 4264 TsaC/Sua5 sequences and 1955 YciO sequences (Supplemental data 1 and 2). To generate the phylogenetic tree, proteins were aligned using MUSCLE (Edgar, 2004). The alignment was trimmed using BMGE (Criscuolo and Gribaldo, 2010) and used to build the maximum likelihood tree using FastTree (Price et al., 2010) with LG+CAT model with bootstrap (1000 replicates) and visualized using iTOL (Letunic and Bork, 2021).

### Sequence similarity networks and gene neighborhood analyses

Sequence Similarity Networks (SSNs) and the corresponding Gene Neighborhood Networks (GNT) were generated using EFI-EST (EFI Enzyme Similarity Tool, <u>efi.igb.illinois.edu/efi-est)</u>(Zallot et al., 2019). Briefly, 54,820 sequences of the PF01300 family between 150 and 350 aa in length were retrieved from Uniprot and subjected to EFI-EST. Each node in the network represents one or multiple sequences that share no less than 70% identity. The initial SSN was generated with an Alignment Score (AS) cutoff set such that each connection (edge) represented a sequence identity above 40%. The nodes of paralogs were colored as given in the legend and visualized using Cytoscape (3.10.1)(Shannon et al., 2003). More SSNs were created by gradually increasing the alignment score cutoff in small increments (usually by 5 AS units). This process was repeated until most clusters were homogeneous in color (AS = 70). The genome neighborhood graphs were generated using Gene Graphics (https://genegraphics.net/)(Harrison et al., 2018).

### Explainable AI methods

To enhance the interpretability of DeepECTF’s multilabel enzyme function predictions, we incorporated an explainable artificial intelligence (XAI) module based on the Local Interpretable Model-agnostic Explanations (LIME) framework (Ribeiro et al., 2016). The approach was specifically adapted to address the multilabel nature of EC number predictions and to provide both local and global insights into the protein sequence segments or residues driving model decisions. Traditional LIME is designed for binary or multiclass outputs, but enzyme function prediction often involves multilabel assignments. To address this, we implemented a multilabel adaptation of LIME, which constructs independent explanation pipelines for each possible label. For each protein sequence input, the XAI module isolates the probability output for a single EC label using the model’s sigmoid activation, enabling LIME to generate label-specific explanations. To optimize computational efficiency, explanations are generated primarily for the label with the highest predicted probability for each input sequence, focusing interpretability efforts on the most relevant predictions.

To balance explanation quality with computational feasibility, local LIME explanations are computed for a representative subsample of up to 500 protein sequences per analysis batch. For each sequence, the module generates a local feature importance map, highlighting which residues in the protein sequence most influenced the model’s prediction for the selected EC label. For proteins outside this subsample, residue-level importance is estimated based on the aggregate statistics from the explained subset. This approach enables the extraction of both local (residue-level) and global (dataset-level) feature importance profiles, which are subsequently exported for downstream analysis. To further dissect model behavior, feature importance scores derived from LIME explanations were stratified by prediction type (correct predictions, paralog errors, non-paralog errors, and repetitions, see Table S1). Each protein sequence is mapped to a prediction category based on expert curation of the model outputs. Residue-level importance values are normalized within each error type, and summary plots are generated to visualize the distribution and magnitude of important sequence segments across different error categories.

The XAI module is fully integrated into the DeepECTF workflow, allowing users to generate explainability outputs alongside standard predictions without additional user intervention (https://github.com/Dias-Lab/XAI_DeepProZyme). All explanation results, including local and global feature importance scores, are made available in standard tabular formats for further interpretation or visualization. This explainable AI approach provides actionable insights into the decision-making process of DeepECTF, supporting both the validation of correct predictions and the systematic investigation of model failures, facilitating the development of more robust and trustworthy protein function prediction systems.

## Results

### Most correctly AI predicted EC numbers are generic or already in the training set

We analyzed 453 *E. coli* proteins functionally annotated in the Kim et al study (Kim et al., 2023) (Table S1)121 had the same EC number in the August 2024 corresponding UniProt annotation(Bateman et al., 2023) and 87% of these were in the version used to generate the training dataset (Fig. 5 and Table S1A) making these an example of training data contamination. 16 proteins had the exact same function labeled in UniProt but without an EC number or with a partial EC number (Fig. 5 and Table S1BC) and are examples of successful annotation propagation (Fig. 1). These 136 cases were all considered correct but not novel predictions (CNN) with confidence scores of 2 (Fig. 5).

**Figure 5.**
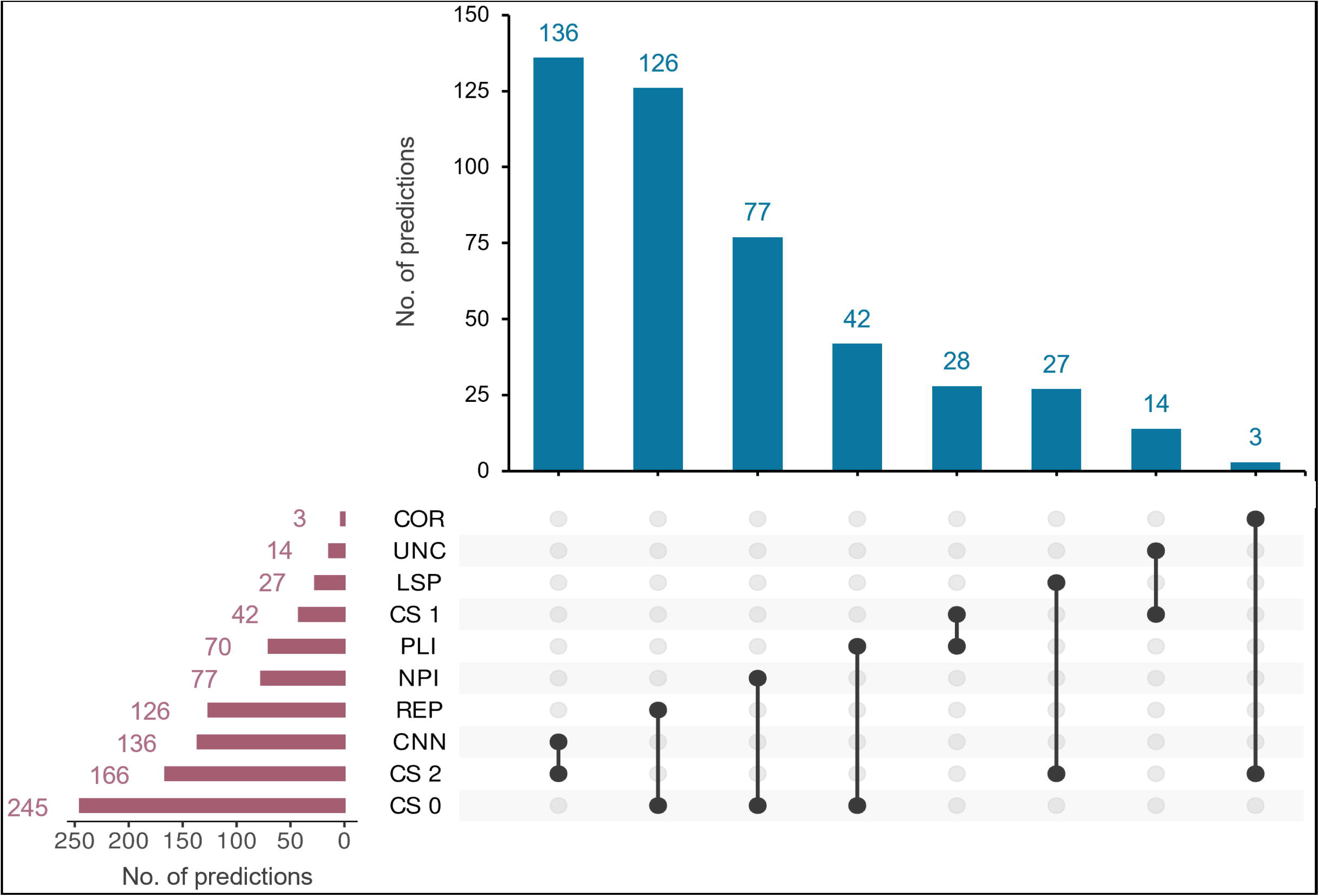
Classification of DeepECTF predictions. The 453 EC number predictions for 453 *E. coli* unknowns were manually classified into categories by comparing the EC number in the UniProt database and labeled with confidence scores from 0 to 2. Data extracted from **Table S1.** Abbreviation: COR, <u>Cor</u>rect prediction; UNC: <u>Unc</u>ertain; LSP, <u>L</u>e<u>s</u>s <u>P</u>recise; PLI, <u>P</u>aralogs incorrect; NPI, <u>N</u>ot-<u>p</u>aralog incorrect; REP, <u>Rep</u>etitions; CNN, <u>C</u>orrect but <u>n</u>ot <u>n</u>ovel; CS, <u>C</u>onfidence <u>s</u>core.

The remaining 317 predictions can be split into proteins with partial or different EC numbers present in the UniProt annotation (63 cases, Table S1B) or with no EC number in the corresponding UniProt entry (254 cases, Table S1C). For 27 predictions, more generic annotations than the existing UniProt annotations were given (less precise cases or LSP in Table S1BC). These were also given a confidence score of 2. The 290 remaining cases were subject to human curation. We combined PaperBlast searches and UniProt and EcoCyc data analyses to group the 290 DeepECTF predictions into different categories with confidence scores from 0 to 2 (Table S1 and Fig. 5). The 42 predictions that could neither be validated nor refuted were labeled as uncertain with a confidence score of 1. Three cases, including YgfF discussed below, were validated by publications not captured in UniProt and can be considered successful predictions (confidence scores of 2). Combining the novel (COR, 3), the obvious (CNN, 136) and the less precise or generic (LSP, 27) predictions brings the number of correct predictions to 166 (36.6%). 245 predictions (54%) were inconsistent with the existing evidence and given confidence scores of 0 (Table S1BC).

### Manual analyses reveal logical inconsistencies in AI predictions of unknowns

The 245 predictions with confidence scores of 0 can be separated into two categories. Those with published evidence refuting the annotation (77 NPI and 42 PLI in Table S1BC) and those that were replications of identical EC numbers (126 REP in Table S1BC). Examples of the first type (n=119) are given in Table 2. For example, YjhQ/b4307 is predicted to be a mycothiol synthase (EC:2.3.1.189), but mycothiol is not a molecule synthesized by *E. coli* and the remaining pathway genes are absent from the genome {BioCyc ID: PWY1G-0). YrhB/b3446 is predicted to be a 6-carboxytetrahydropterin synthase (EC: 4.1.2.50) but *E. coli* already encodes this enzyme (QueD/b2765) and a *queD* mutant lacks this activity (Zallot et al., 2017).

Replication of identical EC numbers occurred for 126 proteins (Fig. 5 and Table S1BC). Repetitions of EC numbers do occur in bacterial genomes, particularly with families that, despite having 4 EC numbers, have generic functions. For example, histidine kinases with different substrate specificities are frequent, with 29 annotated in *E. coli* (Table S1D, top). However, analysis of the protein family domain membership showed that most of the repetitions in these data were errors, except for the less specific annotation cases classified as correct predictions above (Table S1C). For example, out of the 12 proteins annotated as histidine kinase (EC:2.7.13.3), none of them have sequence similarity to histidine kinase families, and eight have been annotated with different and experimentally validated functions (such as ferric enterobactin transport protein FepE for b0587)(Table S1C). For the 15 proteins annotated as “protein-Npi-phosphohistidine-sugar phosphotransferase (EC:2.7.1.69) (or PTS family proteins), 4 were indeed PTS transporters but were given less specific annotations than the ones in UniProt (Table S1C). The 11 remaining were part of transporter families not related to PTS (Table S1B). This type of error may be due to inherent limitations in how AI methods operate. Indeed, if the input features in the training data lack the biological structure and cannot leverage information to distinguish between different functions, the model is expected to make frequency-dependent predictions that reflect the training data, as demonstrated with histidine kinases.

### Correct separation of paralogous group can confirm or refute AI based functional predictions

Many of the incorrect or uncertain (confidence scores of 0 or 1) were part of protein families with paralogs (Fig. 5 and Table S1BC). For example, b2100 was annotated as a dehydro-2-deoxygluconokinase (EC 2.7.1.92 using DeepECTF and as an uncharacterized sugar kinase YegV (EC 2.7.1.-) in UniProt (Table S1B). These two predictions differ by the 4^th^ or last position of the EC number that specifies substrate specificity. This protein is a member of a superfamily of sugar kinases with multiple non-isofunctional paralogous subgroups that phosphorylate different substrates (Table S1D, bottom). The dehydro-2-deoxygluconokinase (EC 2.7.1.92) activity is encoded by another member of this superfamily KdgK/b3526 (Table S1D, bottom). Here, DeepECTF predicted correctly the first 3 digits of the EC number but not the last, making an over-propagation mistake (error 6 in Table 1 and Fig. 1). Correctly separating non-isofunctional paralogous subgroups in a superfamily is difficult, requiring extensive examination of all of the evidence and oftentimes experiments.

We performed an additional, in-depth analysis of the three proteins described using *in vitro* assays in the Kim *et al*. study (YgfF, YdjM, and YciO)(Kim et al., 2023) and show that *in vitro* functionality does not always correspond with *in vivo* functionality.

#### YggF analysis

YgfF is a member of the large Short-Chain Dehydrogenase/Reductase (SDR) superfamily (IPR002347). The Oppermann and Persson groups developed a nomenclature system and HMM-based classification (http://www.sdr-enzymes.org/) that distinguish different functional subgroups of the SDR superfamily(Persson et al., 2009; Kallberg et al., 2010). This resource predicts YgfF is part of the SDR63C/Glucose 1-dehydrogenase subgroup, the activity predicted and validated in the.(Kim et al., 2023) study. This prediction demonstrates the accurate propagation of functional annotation and is a successful prediction.

#### YjdM analysis

YjdM was predicted and shown to catalyze phosphonoacetate hydrolase (EC:3.11.1.2) activity *in vitro*. In *E. coli*, the *yjdM* gene is located upstream of the methylphosphonate catabolism operon (*phnCDEFGHIJKLMNOP)* (Fig. 6). Underscoring the challenges in annotations, the initial report that the protein was involved in phosphonate catabolism was later refuted with additional genetic analyses (Metcalf and Wanner, 1993). The experimentally validated phosphonoacetate hydrolase (PhnA) is part of a non-homologous family and expression of members of this family in *E. coli* suggested that phosphonoacetate hydrolase activity was not present in this organism (Kulakova et al., 1997). Genome neighborhoods of *phnA* show strong clustering with genes encoding phosphonoacetate transporters, phosphonoacetate sensing regulators, and in some cases, enzymes involved in 2-aminoethylphosphonate catabolism (Kulakova et al., 2001). However, except for *E. coli, yjdM* genes are generally not close to phosphonate catabolism or transport genes (Fig. 6). In conclusion, the phosphonoacetate hydrolase activity observed *in vitro* is not supported, and additional *in vivo* experiments are required to confirm the biological role of this enzyme. This prediction was given a confidence score of 1

**Figure 6.**
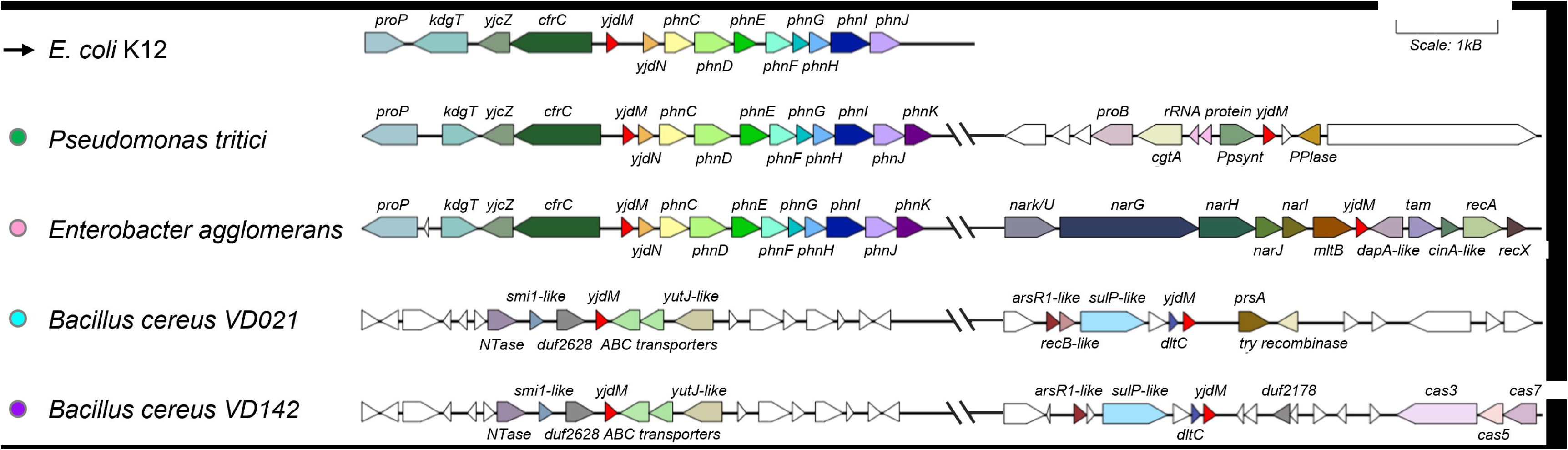
Gene neighborhoods and metabolic reconstructions do not link YdjM to phosphonate degradation. Most *yjdM* genes are not near phosphonate degradation operons. A Sequence Similarity Network of 11,986 IPR004624 family members was generated, and the corresponding genomic neighborhood information is available Table S4.

#### YciO analysis

YciO is a member of the same Pfam family (PF01300) as TsaC/Sua5 and DeepEC predicts YCiO has the same function as TsaC/Sua5. TsaC and Sua5 (the latter known in bacteria as TsaC2) are two types of the well characterized L-threonylcarbamoyladenylate synthase (EC 2.7.7.87) that catalyzes the first step in the synthesis of the universal tRNA modified nucleoside N-6-threonylcarbamoyladenosine or t^6^A (TsaC and Sua5 have a common catalytic domain and differ by the presence of an additional domain in Sua5)(Su et al., 2022; Pichard-Kostuch et al., 2023). The function of TsaC/Sua5 was first elucidated in 2009 (El Yacoubi et al., 2009). A structure-based multi-sequence alignment comparing YciO and TsaC/Sua5 shows that the active site residues of TsaC are largely conserved in YciO, suggesting similar catalytic activities (Fig. 7A). YciO catalyzed the synthesis of L-threonylcarbamoyladenylate from ATP, L-threonine, and bicarbonate, *in vitro* (Kim et al., 2023). However, the activity reported (0.14 nM/min TC-AMP production rate) for *E. coli* YciO is more than four orders of magnitude weaker than that of *E. coli* TsaC (2.8 μM/min) at the same enzyme concentration and similar reaction conditions(Swinehart, W. Deutsch, C. Sarachan, K.L. Luthra, A.; de Crécy-Lagard, V. Swairjo, M. Agris, P.F. Iwata-Reuyl, 2019), consistent with the possibility of a missing partner or a different biological substrate for YciO.

**Figure 7.**
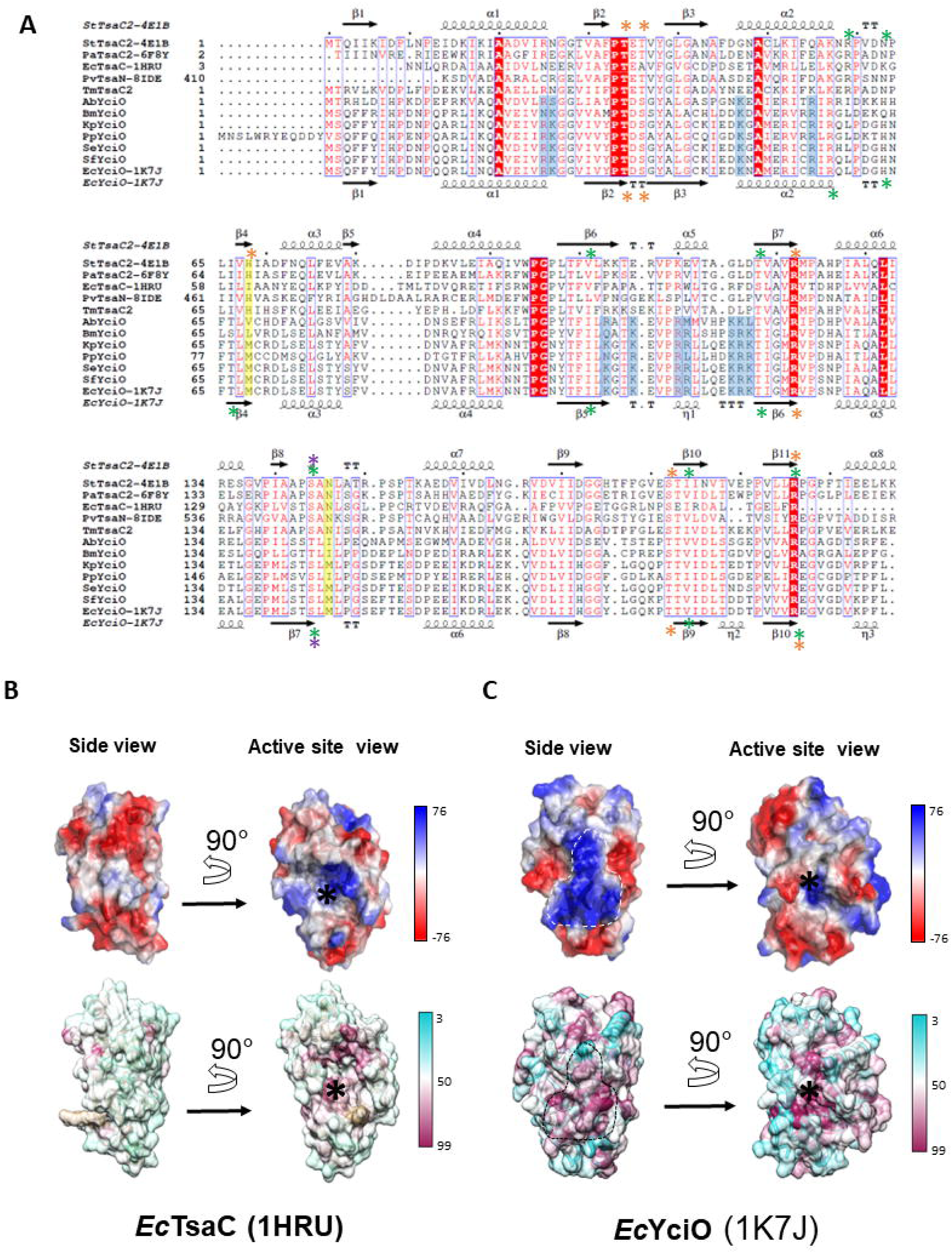
YciO harbors specific molecular surface features. **(A)** Structure-based multi-sequence alignment of TsaC proteins, the TsaC domains of Sua5 proteins and YciO proteins, derived from available crystal structures and AlphaFold structural models. For crystal structures, the PDB IDs are indicated in the sequence name after the hyphen. Secondary structure elements from the crystal structures of *E. coli* TsaC and *E. coli* YciO are displayed above and below the sequences, respectively. YciO-specific conserved basic residues forming the positively charged surface patch of YciO are shaded in blue. Stars above and below the alignment indicate the crystallographically observed substrate binding residues in *St*Sua5, and the corresponding putative substrate binding residues in *Ec*YciO, respectively. Green stars indicate the Mg^2+^-ATP binding residues. Orange stars indicate the binding residues for the L-Threonine substrate. *St*Sua5: *Sulfurisphaera tokodaii* Sua5 (UniProt ID Q9UYB2), *Pa*Sua5: *Pyrococcus abyssi* Sua5 (UniProt Q9UYB2), *Ec*TsaC: *Escherichia coli* TsaC (UniProt P45748), *Pv*TsaN: TsaC domain of Pandoravirus TsaN (UniProt A0A291ATS8). *Tm*TsaC2: *Thermotoga maritima* TsaC2 (by the authors, unpublished, UniProt Q9WZV6), *Ab*YciO: *Actinomycetales bacterium* YciO (UniProt A0A1R4F4R9), *Bm*YciO: *Burkholderia multivorans* YciO (UniProt A0A1B4MSV9), *Kp*YciO: *Klebsiella pneumoniae* YciO (UniProt A6T7X1), *Pp*YciO: *Pseudomonas putida* YciO (UniProt A5W0B1), *Se*YciO: *Salmonella enterica* TciO (UniProt A0A601PQ14), *Sf*YciO: *Shigela flexneri* YciO (UniProt P0AFR6), *Ec*YciO: *Escherichia coli* YciO (UniProt P0AFR4). **(B, C)** Surface representations of the crystal structures of *Ec*TsaC (**B**) and *Ec*YciO (**C**), color-coded by surface electrostatic potential (top), and by positional sequence conservation score calculated from 4264 TsaC/Sua5 sequences and 1955 YciO sequences (bottom). The color keys for both panels are shown on the right. The conserved, YciO-specific, positively charged surface patch (6% of total molecular surface area) is encircled with a dashed line. The active center of TsaC and the putative active center of YciO are marked with asterisks. The figure illustrates that although the active site is conserved in both protein families, the positively charged surface patch is present and conserved only in the YciO family.

YciO does not perform the same function as TsaC/Susa5 *in vivo* experiments(El Yacoubi et al., 2009; Gerdes et al., 2011). Genome neighborhood and structural data suggest that the function of YciO may be related to rRNA rather than tRNA metabolism. The evidence related to rRNA is as follows: 1) the structure of YciO exhibits a large positively charged surface predicted to interact with RNA ((Jia et al., 2002) and Fig. 7B and C). This large positively charged surface is conserved in YciO proteins and is absent in TsaC proteins. 2) in many species, *yciO* genes are colocalized with *rnm* genes (Fig. 8), which encode the recently characterized RNase AM, a 5′ to 3′ exonuclease that matures the 5′ end of all three ribosomal RNAs in *E. coli*(Jain, 2020).

**Figure 8.**
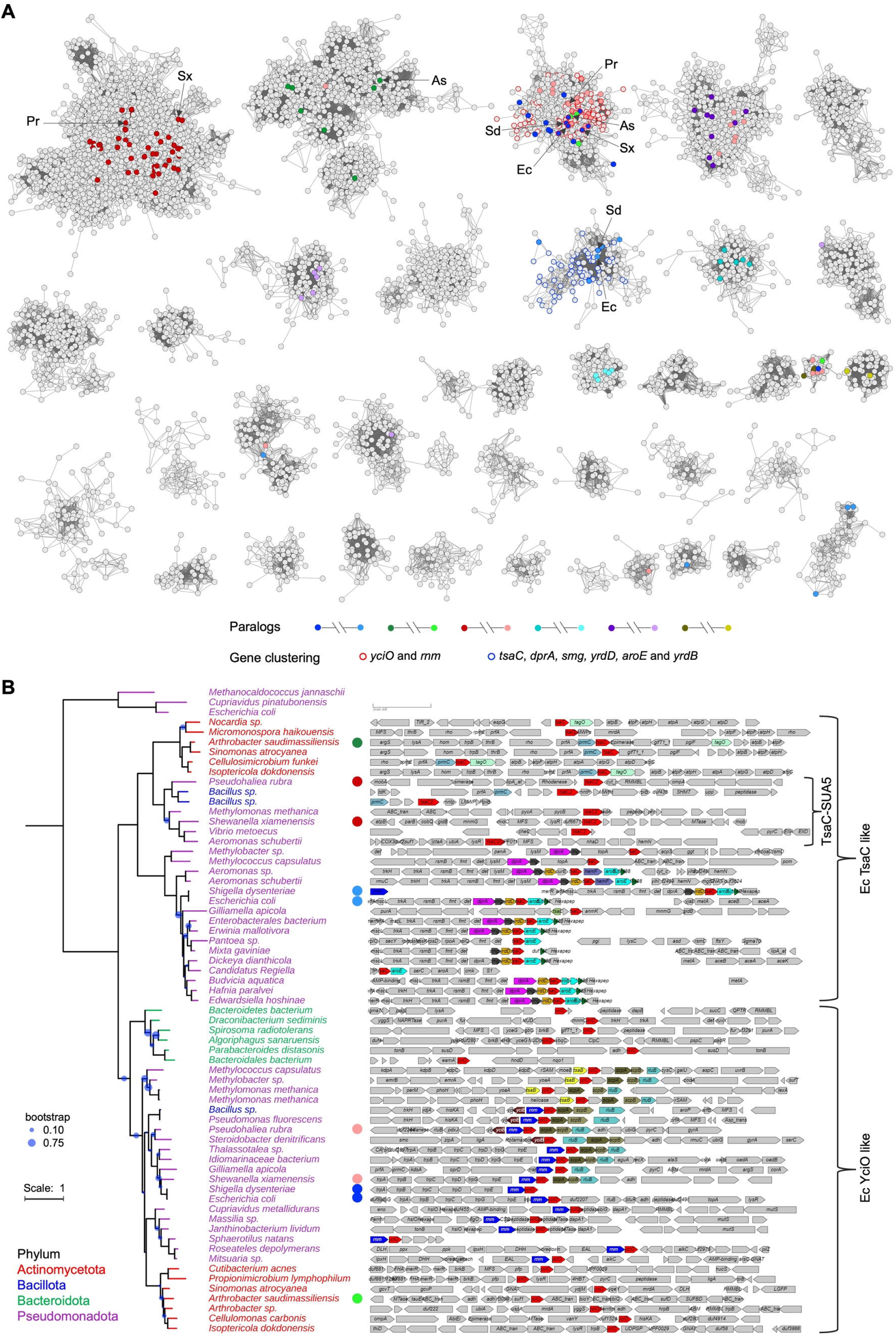
The genes encoding members of the *yciO* and *tsaC* have different genome neighborhood contexts. **(A)** Sequence Similarity Network of 54,820 PF01300 family members between 150 and 350 aa in length generated using EFI-EST. Each node in the network represents one or multiple sequences that share no less than 70% identity. An edge, represented as a line, is drawn between two nodes with alignment scores higher than 70 (similar in magnitude to the negative base-10 logarithm of a BLAST e-value of 1E-70). Paralogs were colored in pairs as indicated. Node borders were colored by gene neighborhood context, *yciO* and *rnm* cluster (red), *tsaC, dprA, smg, yrdD, aroE* and *yrdB* cluster (blue). For better visualization, clusters of less than 40 nodes and singular nodes were hidden. The proteins used in the SSN are available in Table S3. **(B)** Phylogenetic tree of TsaC and YciO proteins. 62 PF01300 proteins were selected from each major cluster in the family SSN and 3 RibB proteins (UniProt Q60364, Q46TZ9, and P0A7J0) were used as the outgroup. Bootstrap values less than 0.75 were indicated by blue dots. Branches and leaves were colored by phyla. TsaC2 proteins that contain YciO/TsaC (PF01300) and Sua5 (PF03481) domains are enclosed in a bracket. The genome neighborhood information is available in Table S4. Selected example organisms were indicated by arrows and a two-letter code in the SSN. Each corresponding genome neighborhood schematic was indicated by a circle in the same color as the node. As, *Arthrobacter saudimassiliensis*; Ec, *Escherichia coli*; Pr, *Pseudohaliea rubra*; Sd, *Shigella dysenteriae*; Sx, *Shewanella xiamenensis*.

Models of enzyme evolution go through promiscuous stages(Ribeiro et al., 2023). TsaC is an enzyme predicted to have been present in the Last Universal Common Ancestor (LUCA)(Gagler et al., 2022; Pichard-Kostuch et al., 2023). YciO is a likely paralog of TsaC (Fig. 7), and as such, it is likely to have residual ancestral catalytic activity that can be detected *in vitro*. In summary, the functional puzzle is far from being solved for proteins of the YciO subgroup, and even if the existing data suggest a role in RNA metabolism, it cannot be the same as TsaC, and the EC number 2.7.7.87 prediction was given a confidence score of 0.

### Feature Importance Patterns identify limitations in the DeepEC model

Application of the XAI module to DeepECTF predictions enabled residue-level analysis of feature importance across protein sequences, stratified by prediction type: correct predictions, paralog errors, non-paralog errors, and repetition errors (Fig. 9). For correct predictions, the feature importance profiles identified a small number of residues-often corresponding to known catalytic or conserved domain positions, showing high contribution scores, indicating that features whose function relies on localized residues for function are more likely to be accurately predicted. In contrast, for all categories of erroneous predictions, feature importance profiles had small deviations from the null values, indicating that the model is not able to leverage information about the protein sequence in making its predictions.

**Figure 9.**
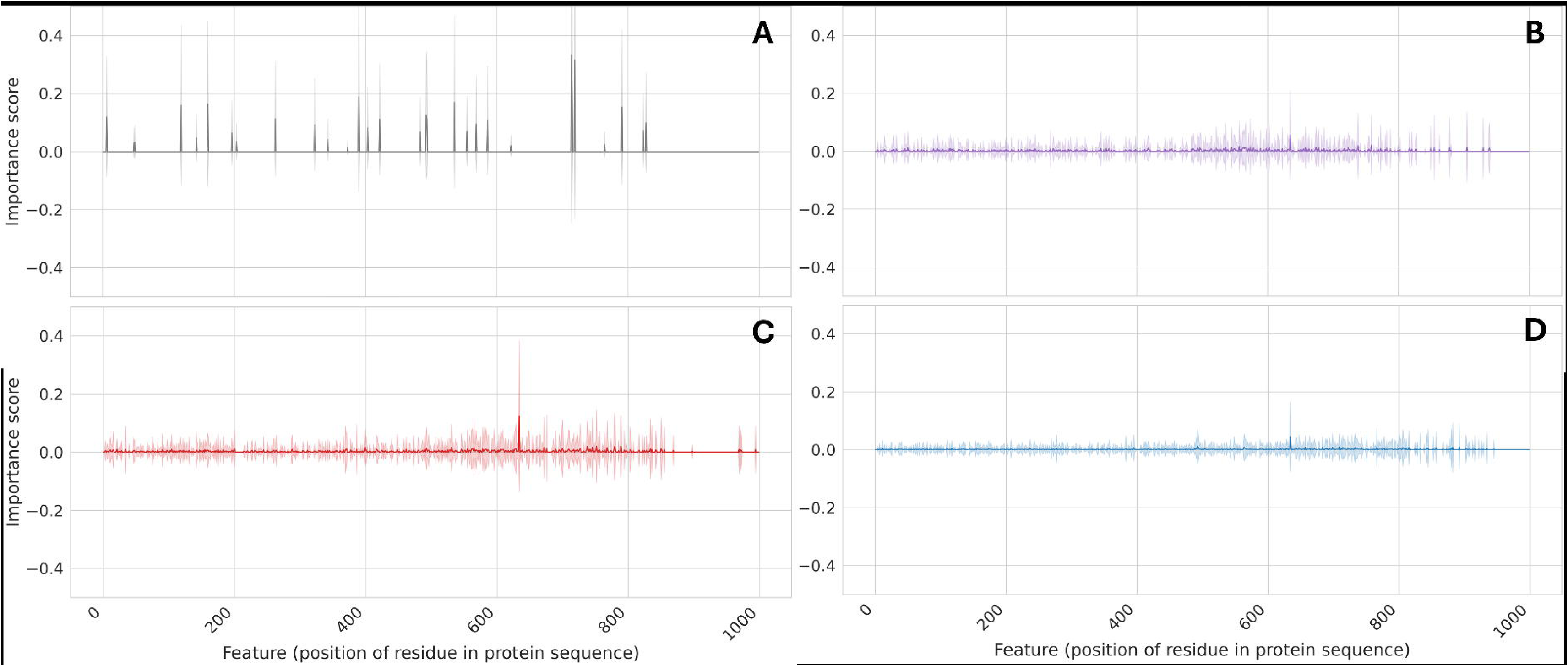
Residue-level feature importance profiles for DeepECTF highlight the distinct interpretability signatures associated with model predictions. Each line plot displays the normalized contribution (average feature importance computed via LIME) of individual residues across protein sequences for four categories: correct predictions (**A**), paralog errors (**B**), non-paralog errors (**C**), and hallucinations or repetitive predictions (**D**). Shaded regions represent standard deviation across instances within each category.

## Discussion

Our expert curation of ML-based EC number predictions for *E. coli* “unknowns” proteins reveals that these methods still have a lot of room to improve. Biases, data imbalance (under- or overrepresented domains, motifs, and functions), generic EC numbers, lack of data structures that enable the inclusion of feature information, architectural limitations (e.g. incapacity of capturing complex patterns, inability to integrate diverse data sources, lack of regularization, etc.), and poor uncertainty calibration all contribute to challenges in training computational models and are likely contributors in frequency dependent predictions (Pucci et al., 2018; Urban et al., 2020; Mardikoraem and Woldring, 2023; Shahbazi et al., 2023). Efforts to improve the quality and consistency of training data and avoid data leakage between training and testing data can help minimize errors but metrics evaluating the uncertainty in predicted protein functions, or an assessment of the likelihood of the label assignment, need to become standard model outputs. In the absence of these metrics, XAI can be used to provide insights into feature importance and model behavior. Many advanced machine learning models, particularly deep neural networks such as DeepECTF, function as ‘black boxes’, where the prediction algorithm is difficult to interpret. Explainable AI (XAI) techniques can help to dissect ‘black-box’ models and understand what input features are driving correct and incorrect predictions (Chennam et al., 2023) and provide insights about which data to include in future models to improve predictions.

Our study also shows that the model’s lack of inclusion of the broader accumulated knowledge in the fields of protein structure and function, biochemical pathways and the absence of a mechanism to identify ‘illogical’ predictions led to erroneous predictions for most proteins of unknown functions, although the model was able to successfully propagate functions among proteins that were included in the training data. We also want to emphasize that *in vitro* activity alone is not sufficient to validate the function of a protein *in vivo* (García-Contreras et al., 2012; Punekar, 2018). Indeed, enzymes evolve by duplication, divergence, and subsequent sub, neo or hypo functionalization (Ribeiro et al., 2023; Birchler, 2025). *In vitro* activities of many enzymes can show promiscuity, which is useful for biotechnological applications (Robinson et al., 2020) but does not guarantee that the protein plays that specific role *in vivo* (Copley, 2015) as shown here with the YciO example. Best practices in functional annotations of enzymes couple biochemical and contextual evidence and reflect the Gene Ontology Consortium definitions, capturing the cellular component as well as the molecular and biological functions (Ashburner et al., 2000).

PLM-driven approaches are becoming mainstream tools for propagating known functional annotations among isofunctional proteins (https://www.uniprot.org/help/ProtNLM). Computational models will be further enabled by the integration of complementary evidence such as structural data to identify active site signature residues (Derry and Altman, 2023; Sajid et al., 2023; Yu et al., 2023), gene neighborhood context (Urhan et al., 2023; Hwang et al., 2024), and chemical reaction specificity (Qian et al., 2024), all of which help to distinguish non-isofunctional paralogous subgroups(Ribeiro et al., 2023). Recent models like MAPred (Rong et al., 2024) align with modern goals of interpretability and transparency in machine learning models (Chennam et al., 2023; Derry and Altman, 2023; Rong et al., 2024) and implements a feature-dropping approach that represents a step toward a more quantitative and nuanced ‘self’ evaluation of model predictions. The future in this space is exciting, even though the limits of these models are just now beginning to be understood.

## Competing interests

The authors declare no competing interests.

## Data availability

Supplemental files are available at FigShare (10.1101/2024.07.01.601547). Table S1 contains the manual review of EC number predictions for 450 *E. coli* unknowns made by DEEP EC. Table S2 contains gene neighborhood information for selected *yjdM* (IPR004624 family) genes shown in Fig. 6. Table S3 includes the identifiers for the PF01300 family members used in Fig. 8 SSN and Table S4 the gene neighborhood information for selected PF01300 family genes. Supplemental data 1 contains the TsaC sequences used to generate positional conservation scores for Fig. 7. Supplemental data 2 contains YciO sequences used to generate the positional conservation scores for Figure 7.

## Acknowledgments

This work was funded by NIGMS (GM110588 to MAS and VDC and GM145937 to IF) and DARPA (HR0012530305 to VDC and RD). We thank Alan J. Bridge (SIB) for critical reading and input on early versions of the manuscript and the peer editors and reviewers who helped us greatly improve the manuscript.

## Contributions

VDC conceived the study, wrote the first draft of the manuscript, and did the analyses for Tables 1-2 and S1. MAS created figure 7 and wrote sections of the manuscript. YY helped generate the data for Table S1-4 and created Figure 1-6 and 8. RD ran the XAI analysis and wrote sections of the manuscript and created Fig. 9. NS ran the mapping and preprocessing of the input data for the XAI analysis and contributed to the implementation of the XAI module in DeepECTF. IF wrote sections of the manuscript. All authors edited the final draft.

## Notes

https://doi.org/10.1101/2024.07.01.601547

### Competing Interest Statement

The authors have declared no competing interest.

### Summary of Updates

The manuscript was substantially revised and additional data was added.

